# Immunogenic shift of arginine metabolism triggers systemic metabolic and immunological reprogramming to prevent HER2+ breast cancer

**DOI:** 10.1101/2024.10.23.619827

**Authors:** Vandana Sharma, Veani Fernando, Xunzhen Zheng, Osama Sweef, Eun-Seok Choi, Venetia Thomas, Saori Furuta

**Author notes:** Contact: Saori Furuta.

## Abstract

Arginine metabolism in tumors is often shunted into the pathway producing pro-tumor and immune suppressive polyamines (PAs), while downmodulating the alternative nitric oxide (NO) synthesis pathway. Aiming to correct arginine metabolism in tumors, arginine deprivation therapy and inhibitors of PA synthesis have been developed. Despite some therapeutic advantages, these approaches have often yielded severe side effects, making it necessary to explore an alternative strategy. We previously reported that supplementing SEP, the endogenous precursor of BH_4_ (the essential NO synthase cofactor), could correct arginine metabolism in tumor cells and tumor-associated macrophages (TAMs) and induce their metabolic and phenotypic reprogramming. We saw that oral SEP treatment effectively suppressed the growth of HER2-positive mammary tumors in animals. SEP also has no reported dose-dependent toxicity in clinical trials for metabolic disorders. In the present study, we report that a long-term use of SEP in animals susceptible to HER2-positive mammary tumors effectively prevented tumor occurrence. These SEP-treated animals had undergone reprogramming of the systemic metabolism and immunity, elevating total T cell counts in the circulation and bone marrow. Given that bone marrow-resident T cells are mostly memory T cells, it is plausible that chronic SEP treatment promoted memory T cell formation, leading to a potent tumor prevention. These findings suggest the possible roles of the SEP/BH_4_/NO axis in promoting memory T cell formation and its potential therapeutic utility for preventing HER2-positive breast cancer.

## Introduction

Arginine is a semi-essential amino acid mainly obtained from outside sources (1) and is mostly metabolized into two opposing pathways: nitric oxide (NO) vs. polyamine (PA) syntheses (2-5). In tumors, arginine tends to be converted to PAs, small polycationic metabolites essential for cell growth and immune suppression (6,7). High levels of PAs help establish the “cold” tumor microenvironment (TME). For example, PAs inhibit formation of cytotoxic (CD8+) memory T cells, compromising immunological responses (7). In addition, tumor-associated macrophages (TAMs), primarily polarized to the immune suppressive M2-type, preferentially produce PAs from arginine, further elevating PA levels in TME (8-10). Such predominance of PA synthesis in tumors is primarily mediated by suppression of the alternative NO synthesis pathway. This is largely due to reduced availability of tetrahydrobiopterin (BH_4_), the essential NO synthase (NOS) cofactor which is often targeted for oxidative inactivation (11). We also showed that pharmacological inhibition of NO in normal mammary glands of wild-type mice induced the formation of precancerous lesions that highly expressed HER2, indicating a potential pathogenic relevance of NO inhibition to HER2+ breast cancer (12). Consistently, by using cell lines of the MCF10A human breast cancer progression series, we showed that cancer progression was closely linked to declines of basal NO and BH_4_ production, along with increases of HER2 and cell proliferation markers (12).

Aiming to correct arginine metabolism in cancer, different strategies have been explored in clinical trials, including arginine deprivation therapy and PA synthesis inhibitor (6,13,14). Despite some therapeutic benefits, these methods have yet yielded serious adverse side effects limiting their usages (15,16). Thus, there is a critical need to develop a less toxic strategy to normalize arginine metabolism in cancer patients. We have been testing the effect of Sepiapterin (SEP), the endogenous BH_4_ precursor, on modulating arginine metabolism in breast tumors and tumor-susceptible individuals and determining its therapeutic efficacy. SEP has so far been extensively utilized for treating metabolic syndrome, such as phenylketonuria, and been reported no dose-limiting toxicity during the Phase I trial (17) unlike other modulators of arginine metabolism (15,16). We previously showed that supplementing SEP to breast cancer cells effectively suppressed PA production, while elevating NO synthesis and also downmodulating HER2 and cell proliferation to the levels similar to normal cells (12,18). We also showed that SEP treatment of the immuno-suppressive M2-type TAMs lowered PA production, while elevating NO production, leading to their functional conversion to the pro-immunogenic M1-type TAMs (18,19). These reprogrammed TAMs, in turn, activated anti-tumor activity of cytotoxic T cells, leading to significant suppression of HER2+ mammary tumor growth.

In the present study, we explored our hypothesis that chronic oral SEP treatment of animals susceptible to HER2+ breast tumors might prevent or prolong the tumor occurrence. We found that long-term oral administration of SEP to a mouse model of HER2-positive mammary tumors strongly prevented tumor formation. As the mechanistic basis of this phenomenon, we observed that SEP induced reprogramming of the systemic metabolism and immunity, elevating total T cell populations in the circulation and bone marrow. Especially, bone marrow-resident T cells are known to be largely memory T cells, indicating the ability of SEP to promote memory T cell formation as part of its tumor preventative capabilities. These findings suggest the therapeutic efficacy of a long-term use of SEP in promoting anti-tumor metabolism and immunity for protection of individuals susceptible to HER2-positive mammary tumors.

## Materials & Methods

### Cell lines

Human monocytic cell line THP–1 (Cat. No. TIB-202™) cells were purchased from American Type Culture Collection (ATCC). MCF10A and CA1d breast cancer cells were obtained from Karmanos Cancer Institute (Detroit, MI) under Material Transfer Agreement (MTA).

### Cell culture & Reagents

THP–1 cells were maintained at a density of 1×10^6^ cells/ml in RPMI 1640 Medium (Thermo Fisher, Waltham, MA, Cat. No. 11835055) supplemented with 10% fetal bovine serum (FBS), 1% Penicillin/Streptomycin, 2 mM GlutaMAX™, 10 mM HEPES buffer, 45 g/L Glucose and 1 mM Sodium Pyruvate (Thermo Fisher, Cat. No. 15140122, Cat. No. 35050061, Cat. No. SH3023701, Cat. No. A2494001 & Cat. No. 11-360-070). CA1d cells were cultured in DMEM/F12 medium with 5 % Horse serum, 1% Penicillin/Streptomycin, Hydrocortisone, Cholera Toxin, and Insulin (Sigma-Aldrich, Inc, St. Louis, MO, Cat. No. H-0888, Cat. No. C8052-2MG & Cat. No. I1882). All the cells were maintained in a 37 °C humidified incubator with 5% CO_2_.

### Antibodies

Here are the antibodies used: primary antibodies for western blot, CD163 (Abcam, Cat. No. ab182422), TNFα (Thermo Fisher, Cat. No. MA523720), and β–Actin (Sigma-Aldrich, Cat. No. A1978); secondary antibodies for western blot: horseradish peroxidase (HRP) conjugated Sheep anti-Mouse IgG (GE Healthcare Life Sciences, Pittsburgh, PA, Cat. No. NA931-1ML), Donkey anti-Rabbit IgG (GE Healthcare Life Sciences, Cat. No. NA934-1ML), and Donkey anti-Goat IgG (ThermoFisher, Cat. No. A16005); primary antibodies for FACS, CD68 (BioLegend, Cat. No. 333821), CD40 (BioLegend, Cat. No. 334305), CD80 (BioLegend, Cat. No. 305205), CD163 (BioLegend, Cat. No. 333609), CD206 (BioLegend, Cat. No. 321109), anti-mouse F4/80 (BioLegend, Cat. No. 123118), anti-mouse CD80 (BioLegend, Cat. No. 104706), anti-mouse CD163 (BioLegend, Cat. No. 155306), and anti-human TNFα (BioLegend, Cat. No. 502943); and for CUT&Tag analysis, anti-H3K27me3 and anti-H3K27Ac antibody (Active Motif 39156, 23254116-11; Active Motif 39135, 23061102-11).

### Modulation of Arginine metabolism

For the induction of NO production, we used SEP, a precursor of NOS cofactor tetrahydrobiopterin: (20 or 100 μM, Career Henan Chemical Co). For NO inhibition, we used the NOS2 inhibitor: 1400W hydrochloride (100 μM, Cayman Chemical, Ann Arbor, MI, Cat. No. 81520). For inhibition of PAs, we used an inhibitor of Arginase I, N-hydroxy-nor-L-arginine (nor-NOHA, 50 μM, Cayman Chemical, Ann Arbor, MI, Cat. No. 10006861).

### In vitro model of TAMs

Human monocytes THP–1 cells were seeded at a density of 3×10^5^ cells/ml and treated with 100 ng/ml phorbol myristate acetate (PMA, InvivoGen, San Diego, CA, Cat. No. tlrl-pma) for 24 hours for their differentiation to nascent (M0) macrophages. M0 cells were then serum starved for 2 hours in X-VIVO™ hematopoietic cell medium (Lonza, Basel, Switzerland, Cat. No. BEBP04-744Q). For M1 polarization (M1-TAMs), M0 cells were treated with PMA (100 ng/ml), 5 ng/ml lipopolysaccharide (LPS, Sigma-Aldrich, Cat. No. L4391-1MG), and 20 ng/ml interferon γ (IFNγ, PeproTech, Cranbury, NJ, Cat. No. 300-02) for 66 hours. For M2 polarization (M2-TAMs), M0 cells were treated with PMA (100 ng/ml), 20 ng/ml interleukin 4 (IL4, PeproTech, Cat. No. 200-04), and 20 ng/ml interleukin 13 (IL13, PeproTech, Cat. No. 200-13) for 66 hours.

### Reprogramming of M2–TAMs to M1-TAMs

THP–1 derived M2–TAMs were treated with 100 μM SEP (Career Henan Chemical Co., Zhengzhou City, China, CAS No. 17094-01-8) every day for 3 days for reprogramming to M1 TAMs. M2–TAMs were also treated with DMSO (Vehicle) and 5 ng/ml LPS and 20 ng/ml IFNγ as negative and positive controls, respectively.

### Immunoblotting

Cell lysates were prepared using the following lysis buffer: 25mM Tris–HCl (pH 8), 150 mM NaCl, 1 mM EDTA (pH 8), 1% NP–40, 5% Glycerol, 1X PhosSTOP (Sigma-Aldrich, Cat. No. 4906837001) and 1X Protease Inhibitor Cocktail (Thermo Fisher, Cat. No. 78425). Total protein concentration of the cell lysates was quantified using the Pierce BCA Protein Assay Kit (Thermo Fisher, Cat. No. 23227). Cell lysates were mixed with the sample buffer (with beta mercaptoethanol) boiled at 100 °C for 10 minutes. Proteins were separated by sodium dodecyl-sulfate polyacrylamide gel electrophoresis (SDS-PAGE). The separated proteins were then electroblotted to methanol–activated polyvinylidene fluoride (PVDF) membranes (Sigma-Aldrich, Cat. No. IPVH00010). Upon transfer, membranes were blocked with 5% non-fat milk in Tris-buffered saline containing 0.1% Tween® 20 (TBST) and incubated overnight with the following primary antibodies: CD163 (Abcam, Cat. No. ab182422), TNFα (Thermo Fisher, Cat. No. MA523720), and β–Actin (Sigma-Aldrich, Cat. No. A1978). Then they were incubated with horseradish peroxidase (HRP) conjugated Sheep anti-Mouse IgG (GE Healthcare Life Sciences, Pittsburgh, PA, Cat. No. NA931-1ML), Donkey anti-Rabbit IgG (GE Healthcare Life Sciences, Cat. No. NA934-1ML) or Donkey anti-Goat IgG (Thermo Fisher, Cat. No. A16005) secondary antibodies (1:5000 dil). Next, the blots were developed with SuperSignal™ West Dura Extended Duration Substrate (Thermo Fisher, Cat. No. 34076) and imaged using Syngene G:BOX F3 gel doc system.

### Flow cytometry (FACS) analysis of cell surface markers

Cells were dissociated from the plates through incubation with PBS containing 5mM EDTA for 15 minutes at 37°C, followed by gentle scraping. Cells were collected into 96 well V bottom plates (USA Scientific, Ocala, FL, Cat. No. 5665-1101), and centrifuged at 1000 rpm for 5 minutes. Cell pellets were washed with Fluorescence-activated cell sorting (FACS) buffer (PBS, 2% FBS) and blocked for 30 minutes on ice in the blocking buffer: 2% FBS, 2% goat serum, 2% rabbit serum and 10 µg/mL human Immunoglobulin G (IgG). Cells were then incubated with fluorochrome-labeled antibodies prepared in FACS buffer for 1 hour on ice. The antibodies used are as follows: CD68 (BioLegend, Cat. No. 333821), CD40 (BioLegend, Cat. No. 334305), CD80 (BioLegend, Cat. No. 305205), CD163 (BioLegend, Cat. No. 333609), CD206 (BioLegend, Cat. No. 321109), anti-mouse F4/80 (BioLegend, Cat. No. 123118), anti-mouse CD80 (BioLegend, Cat. No. 104706), anti-mouse CD163 (BioLegend, Cat. No. 155306), anti-mouse NK1.1 (ThermoFisher, Cat. No. 61-5941-80) and anti-mouse NKp46 (R& D systems, Cat. No. FAB2225F-025). Cells were washed twice with FACS buffer and resuspended in Dulbecco’s phosphate-buffered saline (DPBS) containing 2% formaldehyde. Samples were assayed on the BD FACSCanto™ II system, followed by analyses with FlowJo software.

### FACS analysis of intracellular markers

Cells were treated with 1μg/ml Brefeldin A (Fisher Scientific, Cat. No. B7450) for 5 hours at 37 °C. Upon treatment, cells were collected into 96 well V bottom plates and centrifuged as described above. The samples were then incubated with 1X Fixation/Permeabilization Buffer (R&D systems, Cat. No. FC007) for 12 minutes at 4°C. Following fixation, samples were centrifuged at 1600 rpm for 5 minutes and washed with the following Permeabilization/Washing buffer: PBS, 2% FBS and 0.1% Triton X-100. Samples were then blocked with blocking buffer containing 0.1% Triton X-100 for 10 minutes. After blocking, samples were incubated with fluorochrome-labeled antibodies prepared in permeabilization/washing buffer for 45 minutes at 4 °C. The following antibodies were used: Anti-human TNFα (BioLegend, Cat. No. 502943). Following antibody incubation, samples were washed with permeabilization/washing buffer, resuspended in DPBS containing 2% formaldehyde and assayed on the BD FACSCanto™ II system, followed by analyses with FlowJo software.

### Measurement of BH_4_ production

Cells were washed with ice cold PBS to remove remaining media. Upon washing, cells were scraped gently, and the cell pellets were collected to Eppendorf™ tubes. The cell pellets were vortexed for 10 seconds and flash frozen in liquid nitrogen (N_2_). Then the pellets were thawed at RT. The process was repeated 5 times. Then the pellets were centrifuged at 1000 rpm for 5 minutes, and the supernatants were collected into fresh tubes. The BH_4_ level in cell lysate was measured using an enzyme–linked immunosorbent assay (ELISA) Kit (Abbexa, Sugar Land, TX, Cat. No. abx354211) following manufacturer’s protocol. Cellular BH_4_ levels were normalized using the total protein concentration in cell lysates.

### Animal Study

All *in vivo* experiments were performed in compliance to The Guide for the Care and Use of Laboratory Animals (National Research Council, National Academy Press, Washington, D.C., 2010) and with the approval of the Institutional Animal Care and Use Committee of the University of Toledo, Toledo, OH (Protocol No: 108658) and Case Western Reserve University, Cleveland, OH (Protocol No. 2022-0080). For this study, we only used female mice as animal models of breast cancer which predominantly affects females. (Besides, spontaneous mammary tumor growth in the animal model we used, MMTV-neu/FVB, is only seen for females.) For tumor prevention study, four weeks old female MMTV-neu/FVB (unactivated) (n=20) mice were obtained from the Jackson Laboratory (ID. IMSR_JAX:002376, Bar Harhor, MN, USA), housed under regular conditions and given ad libitum access to acidified water and regular chow. For drug treatment, animals were assigned to experimental groups using simple randomization. They were divided into vehicle (DMSO) vs. SEP (1 mg/kg) treatment group. The drugs were dissolved in acidified drinking water and administered to mice ad libitum starting at 5 weeks of age for 8 months. Tumor incidence was monitored, and the percentage of tumor-free mice was quantified. The body weight, tumor incidence, latency and size were monitored twice a week, and urine and fecal samples were collected once every three weeks. All measurements were performed blinded, and data analysis were unaware of the nature of treatments. If animals met early termination criteria (e.g., morbidity and tumor size>1500 mm^3^), they were euthanized. However, for construction of tumor-free survival curves, the data of all animals (total of 40) were included, and there was no attrition for treatments. At the end of treatment, mammary tumors, spleens, livers, bone marrows, and blood were harvested and processed for further analyses.

### PBMC isolation from mouse blood

At the end of tumor prevention experiment, MMTV-neu/FVB mice mice were grouped into four groups (DMSO with tumors, DMSO without tumors, SEP with tumors, and SEP without tumors). Whole blood was collected by cardiac pucture into EDTA-coated collection tubes, pooled by group, and mixed with the equal volume of PBS-EDTA solution (Thermo Fisher, Cat. No. J60893.K3). To isolate mononuclear cells, the diluted blood was added on top of Lymphoprep medium (STEMCELL technologies, Cat. No. 07851) within SepMate-15 centrifuge tube (STEMCELL technologies, Cat. No. 85415) and spun at 1200 x g for 10 minutes at room temperature for density gradient centrifugation. Mononuclear cells accumulated at the interface between the top serum and bottom Lymphoprep layers were carefully collected and transferred into a separate centrifuge tube. Cells were washed in PBS plus 2% FBS twice, resuspended in RPMI-1640 medium, and cryopreserved until single cell sequencing.

### Single cell sequencing

Single cell sequencing was performed with GEM-X Chromium Single Cell Gene Expression (3′ GEX V3.1) chips on 10X genomics Chromium X Processor at the Discovery Lab in the Global Center for Immunotherapy and Precision Immuno-Oncology at Lerner Research Institute, Cleveland Clinic. Fastq files were mapped to the GRCh38 reference human genome using Cellranger (v5.0.0) (20). Cells containing less than 600 genes and/or more than 30% mitochondrial and ribosomal genes were removed. Sample-specific Seurat objects were created using Seurat (v4.3.0) (21), then normalized using Seurat’s SCTransform method. Samples were integrated based on variable features using Seurat’s IntegrateData function. To help predict cell types, Seurat object was uploaded to BioTuring BbrowserX (22), and cells were annotated using their deep learning-based mouse cell type prediction model (Sub-cell type (version.2) model). BBrowserX was also used to create UMAPs, differentially expressed gene sets, and Chord diagrams.

### Metabolite sample preparation for Metabolomic analysis

For the metabolomic analysis of cells, 1-2 x10^6^ cells each were frozen in freezing solution (90% FBS + 10% DMSO) and stored at − 80 °C. For the metabolomic analysis of plasma, the whole blood was collected from mice into EDTA-treated tubes. Blood was centrifugated at 2000 g for 20 min at 4 °C and then plasma was recovered as the supernatant and stored at − 80 °C. Cell and plasma samples were subjected to untargeted metabolomics analysis at Metabolon, Inc. (Durham, NC, USA).

Analytes were prepared using the automated MicroLab STAR® system (Hamilton Company, Reno, NV, USA). Briefly, samples were thawed and deproteinized by the addition of 4 fold volume of precooled (dry ice) 80% (v/v) methanol extraction solvent containing recovery standard compounds (23). To remove protein, the mixtures were vortexed, incubated with vigorous shaking for 2 min on Glen Mills GenoGrinder 2000 and centrifuged at 15,000 g at for 30 min 4 °C, and the supernatants were collected. The resulting extract was divided into five fractions: two for analysis by two separate reverse phase (RP)/UPLC-MS/MS methods with positive ion mode electrospray ionization (ESI), one for analysis by RP/UPLC-MS/MS with negative ion mode ESI, one for analysis by HILIC/UPLC-MS/MS with negative ion mode ESI, and one sample was reserved for backup. Samples were placed briefly on a TurboVap® (Zymark) to evaporate the organic solvent. The sample extracts were stored overnight under nitrogen before preparation for analysis.

### Metabolomic analysis

Metabolomic analysis was performed on Ultrahigh Performance Liquid Chromatography-Tandem Mass Spectroscopy (UPLC-MS/MS). All methods utilized a Waters ACQUITY ultra-performance liquid chromatography (UPLC) and a Thermo Scientific Q-Exactive high resolution/accurate mass spectrometer interfaced with a heated electrospray ionization (HESI-II) source and Orbitrap mass analyzer operated at 35,000 mass resolution.

The sample extract was dried then reconstituted in solvents compatible to each of the following four methods. Each reconstitution solvent contained a series of standards at fixed concentrations to ensure injection and chromatographic consistency. The first aliquot was analyzed using acidic positive ion conditions, chromatographically optimized for more hydrophilic compounds. In this method, the extract was gradient eluted from a C18 column (Waters UPLC BEH C18-2.1x100 mm, 1.7 μm) using water and methanol, containing 0.05% perfluoropentanoic acid (PFPA) and 0.1% formic acid (FA). The second aliquot was also analyzed using acidic positive ion conditions, however it was chromatographically optimized for more hydrophobic compounds. In this method, the extract was gradient eluted from the same aforementioned C18 column using methanol, acetonitrile, water, 0.05% PFPA and 0.01% FA and was operated at an overall higher organic content. The third aliquot was analyzed using basic negative ion optimized conditions using a separate dedicated C18 column. The basic extracts were gradient eluted from the column using methanol and water, however with 6.5 mM Ammonium Bicarbonate at pH 8. The fourth aliquot was analyzed via negative ionization following elution from a HILIC column (Waters UPLC BEH Amide 2.1x150 mm, 1.7 μm) using a gradient consisting of water and acetonitrile with 10 mM Ammonium Formate, pH 10.8. The MS analysis alternated between MS and data-dependent MSn scans using dynamic exclusion. The scan range varied slighted between methods but covered 70-1000 m/z. Raw data files are archived and extracted as described below.

### Metabolomic data analysis

The bioinformatics system consisted of four major components: the Laboratory Information Management System (LIMS), the data extraction and peak-identification software, data processing tools for QC and compound identification, and a collection of information interpretation and visualization tools for use by data analysts. Metabolon LIMS system enables fully auditable laboratory automation that encompasses sample accessioning, sample preparation and instrumental analysis and reporting and advanced data analysis. All of the subsequent software systems are grounded in the LIMS data structures. The hardware and software foundations for these informatics components were the LAN backbone, and a database server running Oracle 10.2.0.1 Enterprise Edition.

Raw data was extracted, peak-identified and QC processed using Metabolon’s hardware and software built on a web-service platform utilizing Microsoft’s .NET technologies. Compounds were identified by comparison to library entries of purified standards or recurrent unknown entities. Metabolon maintains a library based on authenticated standards that contains the retention time/index (RI), mass to charge ratio (m/z), and chromatographic data (including MS/MS spectral data) on all molecules present in the library. Biochemical identifications are based on retention index within a narrow RI window of the proposed identification, accurate mass match to the library +/- 10 ppm, and the MS/MS forward and reverse scores between the experimental data and authentic standards. The MS/MS scores are based on a comparison of the ions present in the experimental spectrum to the ions present in the library spectrum. The use of all three data points can be utilized to distinguish and differentiate biochemicals. More than 3300 commercially available purified standard compounds have been registered into LIMS for analysis on all platforms. Additional mass spectral entries have been created for structurally unnamed biochemicals which have the potential to be identified by future acquisitions.

A variety of curation procedures were carried out to ensure that a high quality data set was made available for statistical analysis and data interpretation. The QC and curation processes were designed to ensure accurate and consistent identification of true chemical entities, and to remove those representing system artifacts, mis-assignments, and background noise.

Peaks were quantified using area-under-the-curve. For studies spanning multiple days, a data normalization step was performed to correct variation resulting from instrument inter-day tuning differences. Essentially, each compound was corrected in run-day blocks by registering the medians to equal one (1.00) and normalizing each data point proportionately (termed the “block correction”). For studies that did not require more than one day of analysis, no normalization is necessary, other than for purposes of data visualization. In certain instances, biochemical data may have been normalized to an additional factor (e.g., cell counts, total protein as determined by Bradford assay, osmolality, etc.) to account for differences in metabolite levels due to differences in the amount of material present in each sample.

Further bioinformatics analyses on curated data, including pathway analyses and ROC curve analyses were performed using Metaboanalyst 6.0 package (McGill University, Montreal, Quebec, Canada) (24). Heatmaps were created using the SRPLOT software, while volcano plots were created using Graphpad Prizm Version 10.5.

### Cut & Tag analysis of bone marrow cells

Bone marrows of drug treated animals were subjected to CUT & Tag analysis by Active Motif (Carlsbad, CA, USA). Analytes were prepared as previously described with modifications (25). Briefly, frozen cell pellets were thawed, and nuclei were isolated and incubated overnight with Concanavalin A beads and 1.3 µl of the primary anti-H3K27me3 or anti-H3K27Ac antibody (Active Motif 39156, 23254116-11; Active Motif 39135, 23061102-11) per reaction. After incubation with the secondary anti-rabbit antibody (1:100), beads were washed, and tagmentation was performed at 37℃ using protein-A-Tn5. Tagmentation was halted by the addition of EDTA, SDS and proteinase K after which DNA extraction, and ethanol purification was performed, followed by PCR amplification and barcoding (see Active Motif CUT&Tag kit, Cat. No. 53160 for recommended conditions and indexes). Following SPRI bead cleanup (Beckman Coulter), the resulting DNA libraries were quantified and sequenced on Illumina’s NextSeq 550 (8 million reads, 38 paired end). Reads were aligned using the BWA algorithm (mem mode; default settings) to the mouse genome (mm10) (26). Duplicate reads were removed, and only reads that mapped uniquely (mapping quality >= 1) and as matched pairs were used for further analysis. Alignments were extended in silico at their 3’-ends to a length of 200 bp and assigned to 32-nt bins along the genome. The resulting histograms (genomic “signal maps”) were stored in bigWig files. Peaks were identified using the MACS 3.0.0 algorithm in bedpe mode at a cutoff of q-value 0.05, without control file, and with the –nolambda option. Peaks that were on the ENCODE blacklist of known false ChIPSeq peaks were removed. Signal maps and peak locations were used as input data to Active Motifs proprietary analysis program, which creates Excel tables containing detailed information on sample comparison, peak metrics, peak locations and gene annotations. For differential analysis, reads were counted in all merged peak regions (using Subread), and the replicates for each condition were compared using DESeq2 (27). Other key software used were as follows: bcl2fastq2 (v2.20) (processing of Illumina base-call data and demultiplexing); Samtools (v0.1.19) (processing of BAM files); BEDtools (v2.25.0) (processing of BED files); wigToBigWig (v4) (generation of bigWIG files); and Subread (v1.5.2) (counting of reads in BAM files for DESeq2).

The data (BED files) were visualized by using the custom-track function of UCSC Genome Browser and specifying specific genomic regions of interest. The pathway analyses of differentially expressed gene sets were performed using the Metascape software. Cell analyses based on epigenomic profiles were performed using the Cellkb software (Combinatics, Ltd.).

### Statistical Data Analysis

All the experiments were performed in replicates (n ≥ 3 for *in vitro* analyses and n=7 for animal experiments) ensuring the adequate statistical power (power ≥ 0.8; error ≤ 0.05; and effect size ≥ 0.8, calculated with G*Power) based on our previous studies (28,29). Statistical analyses were performed using Graphpad Prism 10.5, and unless otherwise indicated, two-tailed t-tests, Mann-Whitney U test (non-parametric), or two-way ANOVA test with Bonferroni post hoc test (multi-comparisons)(30) were performed to obtain the statistical significance of the mean difference. *P* values ≤ 0.05 were considered statistically significant. Flow cytometry data analyses were performed using FlowJo Version 10.5.

## Results

### Reprogramming arginine metabolism inhibits HER2+ breast cancer growth

Arginine is a semi-essential amino acid mainly obtained from outside sources (1) and is mostly metabolized into two opposing pathways: NO vs. PA syntheses (**Fig. 1A**)(2-5). In cancer, arginine tends to be converted to PAs, promoting cancer cell growth and immune suppression (6,7). Elevated PA synthesis is largely due to reduced NO synthesis owing to lower availability of tetrahydrobiopterin (BH_4_), the essential NO synthase cofactor (**Fig. 1A**)(12,18). We previously showed that pharmacological inhibition of NO, which would direct arginine metabolism towards PA synthesis, in normal mammary glands of mice induced the formation of precancerous lesions that highly expressed HER2, indicating a pathogenic relevance of NO inhibition to HER2+ breast cancer (12). By using MCF10A human breast cancer progression series (normal MCF10A >> cancerous CA1d), we found that cancer progression of this series was also linked to declines of basal BH_4_ and NO production (**Fig. 1B**) as well as the increases in the levels of HER2 and a proliferation marker Ki67 (12). Conversely, when CA1d cancer cells were treated with SEP (the endogenous BH_4_ precursor) (**Fig. 1C**), BH_4_ and Ki67 levels were normalized to the levels of MCF10A cells (**Fig. 1D, E**)(12). SEP treatment of the progression series lowered PA levels, while elevating NO levels, inducing the shift of arginine metabolism from PA to NO syntheses (18). In addition to arginine metabolism, SEP treatment of CA1d cells normalized the levels of a group of other metabolites to the levels of MCF10A cells, greatly differentiating their levels from those in control CA1d cells (**Fig. 1F, G**). These normalized metabolites largely belonged to fatty acid and nucleotide metabolisms involved in energy production (**Fig. 1H**)(31). The results show that SEP induces reprogramming of arginine metabolism and other metabolic pathways to inhibit HER2+ breast cancer cell growth.

**Figure 1.**
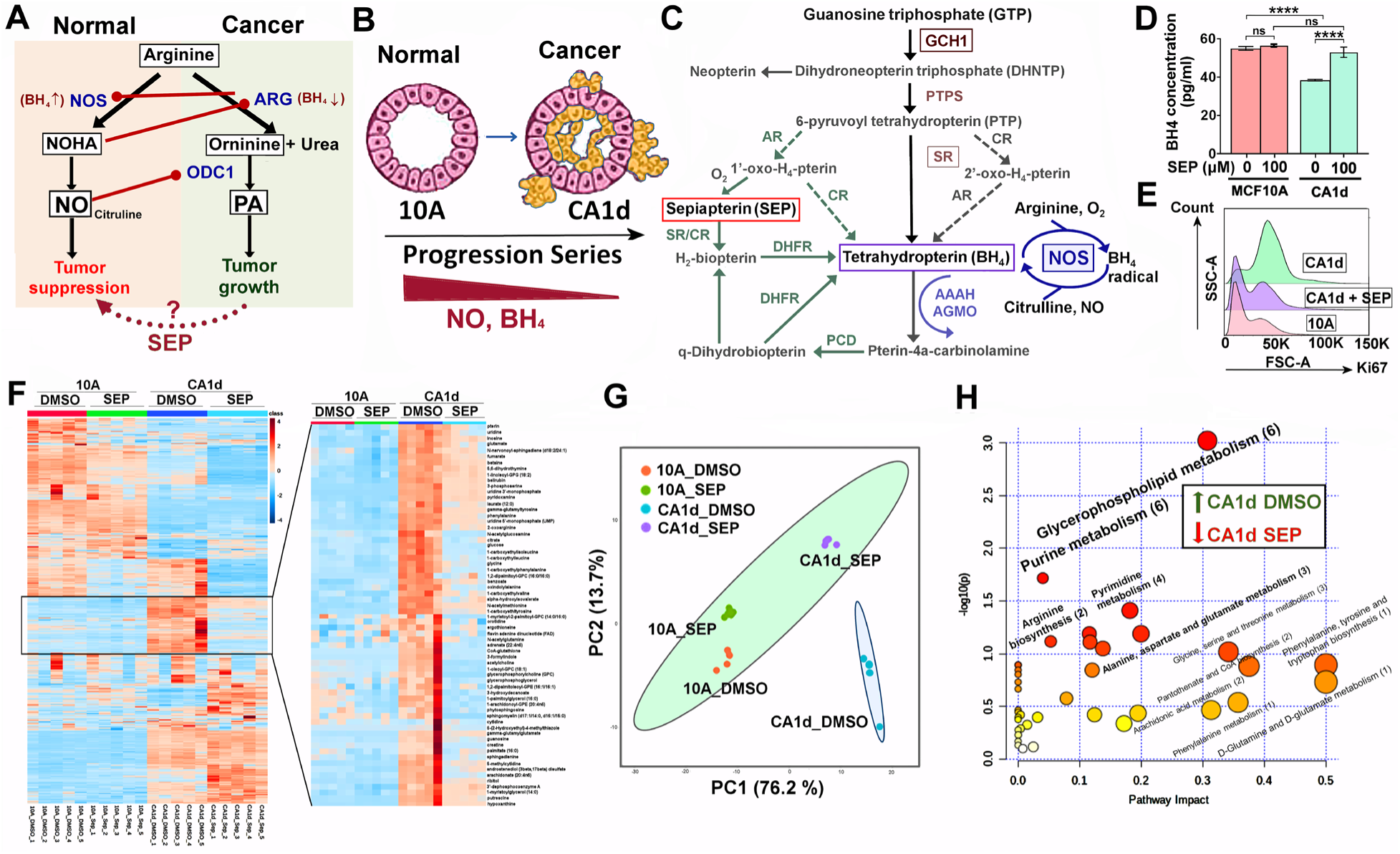
Reprogramming arginine metabolism inhibits the growth of HER2+ breast cancer cells. **A.** Bimodal arginine metabolism leading to two antagonizing pathways: NO vs. PA syntheses. NOS: nitric oxide synthase, NOHA (Nω-hydroxy-nor-arginine); ARG: arginase; ODC1: ornithine decarboxylase 1. **B.** MCF10A (normal) >> CA1d (invasive) in MCF10A human breast cancer progression series showing the decrease of NO and BH_4_ levels along with cancer progression. **C.** BH_4_ biosynthesis pathways featuring sepiapterin (SEP) as a precursor. **D.** Reduced BH_4_ levels in cancerous CA1d compared to normal MCF10A cells, and normalization of the levels after SEP (100 µM) treatment for 2 days (12). **E.** Normalization of a proliferation marker Ki67 levels in CA1d cells treated with SEP. **F.** (Left) Heatmap of metabolite levels in MCF10A cells vs. CA1d cells treated with control DMSO vs. SEP (n=5). (Right) Metabolites normalized in CA1d after SEP treatment. **G.** Principal component analysis (PCA) of metabolites normalized in SEP-treated CA1d. Note the co-localization of SEP-treated CA1d with 10A (DMSO or SEP-treated) and a segregation of DMSO-treated CA1d. **H.** Pathway analysis of metabolites normalized (downmodulated) in SEP-treated CA1d. Note the significant involvements of fatty acid and nucleotide metabolisms in the normalized pathways.

### Redirecting arginine metabolism reprograms tumor-associated macrophages (TAMs)

HER2+ breast tumors have poor immunogenicity due to abundant immune-suppressive cells, including M2-TAMs (10,32,33). TAMs consist of the immune-stimulatory, tumoricidal M1-type and immune-suppressive, pro-tumor M2-type, which could be recapitulated by specific cytokine treatment *in vitro* (**Fig. 2A**). M1 vs. M2 TAM formation largely depends on differential arginine metabolism (18). M1-TAMs convert arginine to NO for pro-inflammatory signaling, whereas M2-TAMs convert arginine to PAs for anti-inflammatory signaling (**Fig. 2B**)(18,34). In fact, inhibition of NO production in M1 macrophages with an NOS2 inhibitor 1400W reduced an M1 marker TNFα level, while elevating an M2 marker CD206 level. Conversely, inhibition of PA production in M2 macrophages with an arginase inhibitor NOHA (Nω-hydroxy-nor-arginine) elevated TNFα, while downmodulating CD206 (**Fig. 2C**). Such differential arginine metabolism in M1 vs M2 TAMs is likewise attributed to the different BH_4_ availability. M2 macrophages produced significantly lower levels of BH_4_ than M1 macrophages. However, SEP treatment of M2 macrophages restored the levels of BH_4_ similar to those of M1 macrophages (**Fig. 2D**). At the same time, SEP treated M2 macrophages came to express an M1 marker TNFα, while dramatically downmodulating an M2 marker CD163 (**Fig. 2E**)(18,19). In addition to arginine metabolic pathways, SEP treatment also lowered a group of metabolites in M2 macrophages to the levels of M1 type, greatly differentiating their levels from those in control M2 macrophages (**Fig. 2F, G**). Such metabolites belonged to the pathways involved in energy production, including TCA cycle, lipid and amino acid metabolism (**Fig. 2H**). These results demonstrate that SEP induces reprogramming of arginine metabolism in M2 macrophages and converts their phenotype to M1 type. In fact, we previously reported that these SEP-treated M2 macrophages are functionally reprogrammed to M1 macrophages and able to induce effector T cells to kill tumor cells (19).

**Figure 2.**
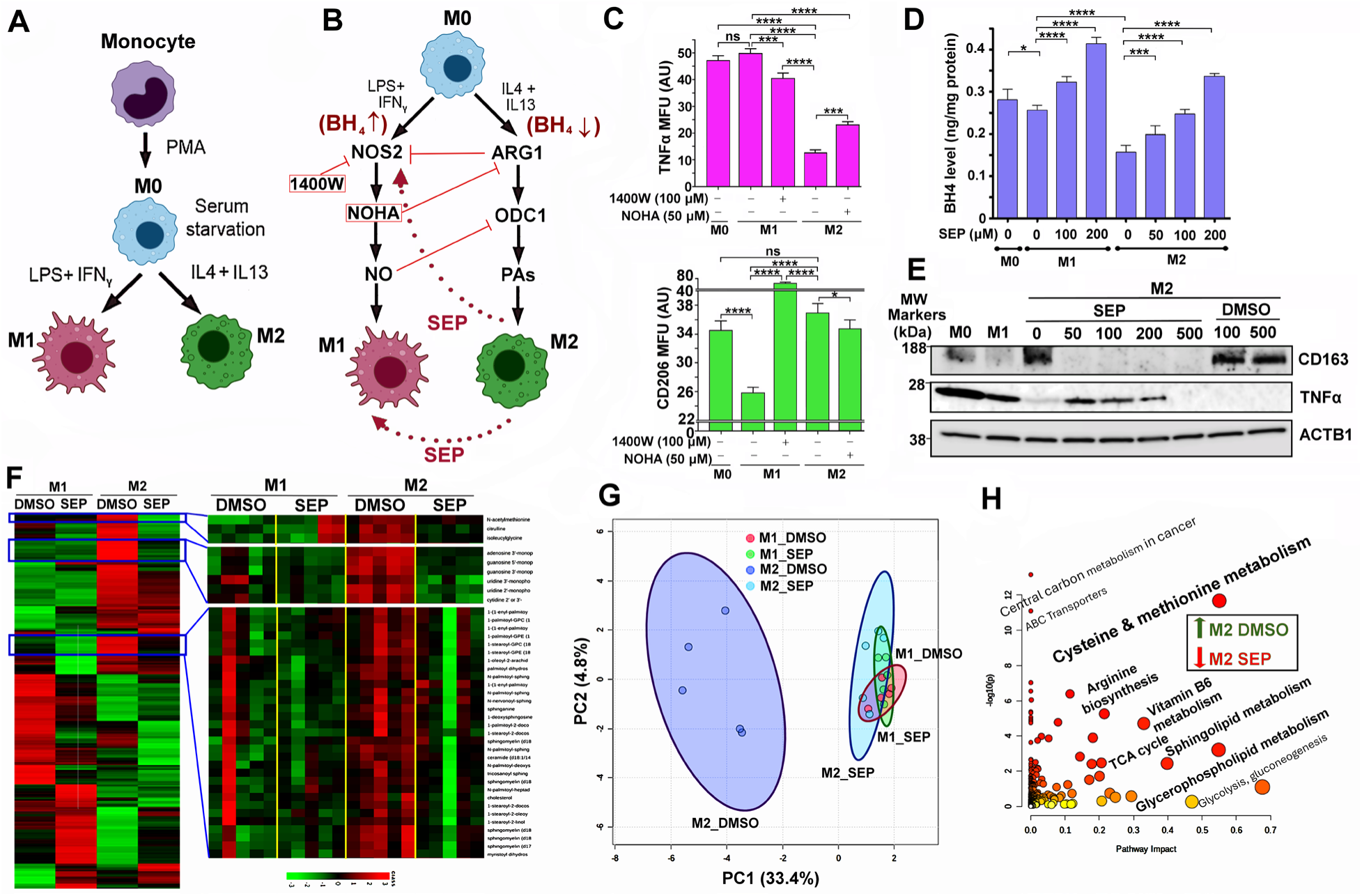
Redirecting arginine metabolism reprograms tumor-associated macrophages (TAMs). **A.** Scheme for differentiation of monocytes into naïve (M0) macrophages and subsequent M1 vs. M2 polarization *in vitro*. **B.** Discrete arginine metabolic pathways in M1 vs. M2 macrophages and their antagonistic relationships. Scheme for phenotypic conversions between M1 and M2 macrophages by inhibiting NOS2 in M1 (M1→M2 conversion) vs. ARG1 in M2 (M2→M1 conversion) or providing SEP to M2 (M2→M1 conversion) (18,19). **C.** (Top) M1 marker TNFα levels in different subsets of macrophages (M0, M1, or M2) treated with an NOS2 inhibitor 1400W or ARG1 inhibitor NOHA. MFU: Mean fluorescence unit. (Bottom) M2 marker CD206 levels in macrophage subsets treated with 1400W or NOHA. Note that 1400W treatment of M1 cells reduced TNFα levels, but increased CD206 levels. Conversely, NOHA treatment of M2 cells elevated TNFα levels, but decreased CD206 levels. **D.** Reduced BH_4_ levels in M2 macrophages, compared to M0 and M1 macrophages, and dose-dependent normalization of the levels after SEP (50, 100, or 200 µM) treatment for 3 days. **E.** Treatment of M2 macrophages with SEP (50, 100, or 200 µM) lowered M2 marker (CD163) levels, while elevating M1 marker (TNFα) levels. Note that the decrease of TNFα levels in M2 cells treated with the extremely high (500 µM) SEP was presumably due to reduced viability of cells. **F.** (Left) Heatmap of metabolite levels in M1 vs. M2 macrophages treated with control DMSO vs. SEP (n=5). (Right) Metabolites normalized in M2 macrophages after SEP treatment. **G.** PCA of metabolites normalized in SEP-treated M2 macrophages. Note the co-localization of SEP-treated M2 with M1 (DMSO or SEP-treated) macrophages and segregation of DMSO-treated M2. **H.** Pathway analysis of metabolites normalized (downmodulated) in SEP-treated M2 macrophages. Note the significant involvements of these metabolites in energy production, including TCA cycle, lipid and amino acid metabolism.

### Long-term SEP treatment prevents mammary tumor occurrence of MMTV-neu mice, while inducing systemic immunological reprogramming

The above results showed that SEP could normalize the phenotypes of both breast cancer cells and TAMs. We then tested whether SEP could suppress tumor formation *in vivo*. We gave SEP (1 mg/kg) in drinking water to MMTV-neu mice starting at their prepubertal stage (5 weeks old) for 8 months. These mice were expected to develop spontaneous single-focal mammary tumors at the latencies of 6-12 months (19). While 90% of DMSO-treated mice had developed tumors within 8 months, over 50% of SEP-treated mice were completely protected from tumor occurrence (**Fig. 3A**).

**Figure 3.**
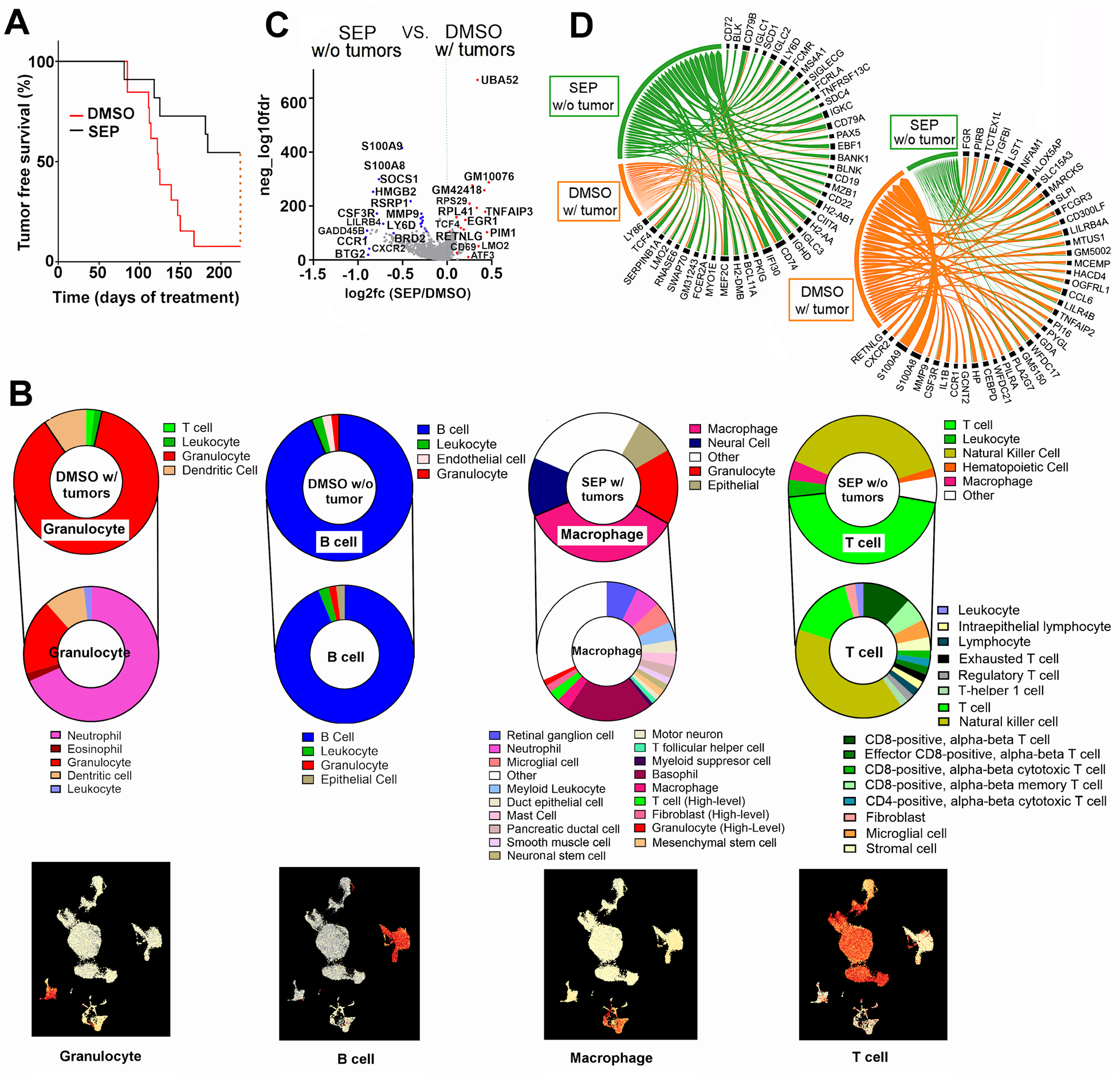
Long-term oral SEP treatment prevents mammary tumor occurrence of MMTV-neu mice, while inducing systemic immunological reprogramming. **A.** Tumor-free survival curves of MMTV-neu mice treated with control DMSO vs. SEP (1 mg/kg, n=20) in drinking water. **B.** Major cell types (top) and sub cell types (middle) determined by single cell sequencing of PBMCs of DMSO with tumor, DMSO without tumor, SEP with tumor, and SEP without tumor groups. (Bottom) UMAPs featuring the major cell type for each group. Note the enrichment of T cells in PBMCs of SEP without tumor group. **C.** Volcano plot for differentially expressed genes in PBMCs of MMTV-neu mice which were SEP-treated and did not develop tumors (SEP without tumors) vs. DMSO-treated and developed tumors (DMSO with tumors). **D.** Chord diagrams of genes elevated (left) or downmodulated (right) in PBMCs of SEP without tumor group compared to those of DMSO with tumor group

To explore the mechanism of such a strong tumor-preventative effect of SEP, we performed single cell sequencing analysis on PBMCs of these treated animals. DMSO- or SEP-treated animals were composed of 4 groups: 1) DMSO with tumors (90% DMSO mice); 2) DMSO without tumors (10% DMSO mice); 3) SEP without tumors (55% SEP mice); and 4) SEP with tumors (45% SEP mice) (**Fig. 3B**). Cell type analysis showed that PBMCs of SEP without tumor group were mainly T cells (CD8+ and CD4+) and NK cells. PBMCs of DMSO without tumor group were mostly B lymphocytes, attesting to the general anti-tumor roles of lymphocytes (35). Conversely, PBMCs of DMSO with tumor group were largely granulocytes, especially, neutrophils, while those of SEP with tumor group were mostly macrophages, attesting to pro-tumor roles of myeloid cells (**Fig. 3B**)(36).

For further analyses, we focused on comparisons between SEP without tumor vs. DMSO with tumor groups. Differentially expressed gene set analysis on PBMCs of both groups showed that SEP without tumor group elevated a number of pro-immunogenic genes involved in T /NK cell functions (TNFAP3, PIM1, LMO2, ATF3, CD69, CD74, IF130, TCF4, and MEF2C)(37-45), while lowering a list of immune suppressive genes (S100A8/9, MMP9, TGFB1, LILR4B, LILRB4A, CCR1, HP, CSF3R, and SOCS1) (**Fig. 3C, D**)(46-53). These results suggest that anti-tumor effects of SEP were linked to the systemic immunological reprogramming that elevated anti-tumor lymphocytes.

### Long-term SEP treatment induces systemic metabolic reprogramming in MMTV-neu mice

To examine the potential causes for the different immunological landscapes between SEP-treated (no tumor) and DMSO-treated (with tumor) groups, we compared their systemic metabolic profiles by analyzing their plasma metabolites. We saw global decreases of metabolites in SEP-treated group compared to DMSO-treated group (**Fig. 4A**). Analysis of differentially represented metabolites revealed that those downmodulated in SEP-treated group mostly belonged to energy production and immune suppression pathways, including TCA cycle, nucleotide, phenylalanine, and tryptophan metabolism (**Fig. 4B**)(54-56). Such metabolites downmodulated in SEP-treated group included N-acetylneuraminate, N-acetylalanine, and taurodeoxycholate (**Fig. 4C**) involved in immunosuppression/exhaustion as well as carcinogenesis (57-62). Conversely, metabolites upregulated in SEP-treated group mainly belonged to pro-inflammatory pathways, including histidine/histamine, taurine, and CoA metabolism (**Fig. 4B**)(63-65). Those metabolites elevated in SEP-treated group included linolenate (α and γ), S-acetylcisteine, citrate, and succinoyltaurine (**Fig. 4C**), involved in immune activation and tumor suppression (66-72).

**Figure 4.**
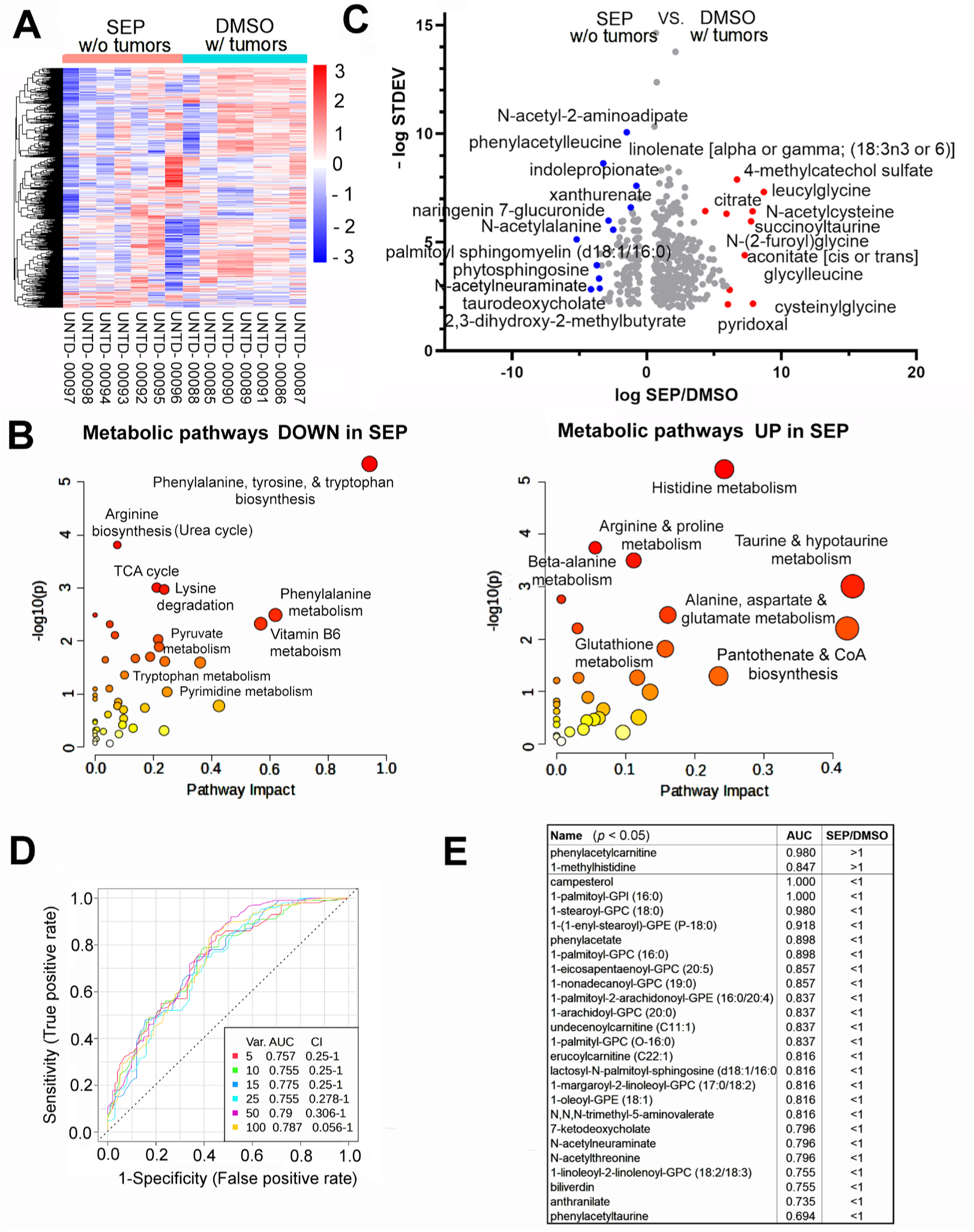
Long-term oral SEP induces systemic metabolic reprogramming in MMTV-neu mice. **A.** Heatmap of metabolite levels in the plasma of SEP without tumor group vs. DMSO with tumor group of MMTV-neu mice (n=7). Note the global decreases of metabolites in SEP without tumor group. **B.** Metabolic pathways downregualted (left) and upregulated (right) in SEP without tumor group compared to DMSO with tumor group. Note that metabolites lowered in the former group mostly belonged to energy production and immune suppression pathways, including TCA cycle, nucleotide, phenylalanine, and tryptophan metabolism. Conversely, metabolites elevated in the former group mainly belonged to pro-inflammatory pathways, including histidine/histamine, taurine, and CoA metabolism. **C.** Volcano plot for differentially represented metabolites in the plasma of SEP without tumor group vs. DMSO with tumor group. Note the significant reduction of tumor-associated metabolites, including N-acetylneuraminate and palmotyl sphingomyelin (59,102). Conversely, the increases in citrate and aconitate indicate inhibition of the TCA cycle. **D.** Receiver operating characteristic (ROC) curve analysis (multivariate, random forest regression) on differentially represented metabolites. **E.** Metabolites with the largest Area under Curve values (AUC∼1) and statistical sifnicance (*p*<0.05).

To determine potential biomarkers, we analyzed Receiver Operating Characteristic (ROC) curves (multivariate, random forest regression) (**Fig. 4D**). The analysis identified a list of metabolites differentially represented in SEP-treated vs. DMSO-treated groups with the highest accuracy (i.e., Area under Curve values (AUC)∼1). Metabolites with the highest accuracy (AUC=1.0-0.98) were campesterol, 1-palmitoyl-glycerophosphoinositol (GPI)(LysoPI), and 1-stearoyl-GPC (LycoPC) which were all involved in immunosuppression (73-75) and downmodulated in SEP-treated group (**Fig. 4E**). Conversely, the metabolite elevated in SEP-treated group with the highest accuracy (AUC=0.98) was phenylacetylcarnitine produced by gut microbial metabolism (**Fig. 4E**)(76,77), indicating SEP’s ability to reprogram the metabolism of gut microbiota in addition to the host. Another metabolite elevated in SEP-treated group with high accuracy (AUC=0.847) was 1-methylhistidine, which is a marker for muscle turnover during extensive exercise and is also an antioxidant (78,79). This could be due to high levels of NO production after SEP treatment promoting muscle development and functions (80). These results altogether demonstrate that SEP-treatment induced systemic metabolic reprogramming to promote immunological responses.

### Bone marrow of SEP-treated MMTV-neu mice has increased levels of total T cells

To further examine the systemic immunological reprogramming by SEP-treatment, we determined the epigenetic profiles of the bone marrows (BM) of drug-treated mice by CUT&Tag analysis for H3K27me3 (suppression) and H3K27ac (activation) histone marks. The heatmaps and peak sizes of the epigenome showed that both histone marks were higher in SEP without tumor group than DMSO with tumor group (**Fig. 5A, B**). A list of genes with elevated H3K27me3 marks (more strongly suppressed) after SEP treatment mostly belonged to pathways involved in immune suppression (e.g., lymphocyte apoptosis and IL5 production)(**Fig. 5C, left**) (81). Conversely, a gene set with elevated H3K27ac mark (more strongly activated) after SEP treatment largely belonged to pathways involved in pro-inflammatory responses (e.g., leukocyte activation and IL-6 production)(**Fig. 5C, right**)(82). Such SEP-activated genes included those involved in T cell activation and development (IL6RA, IL7R, IFNGR2, and CD44) (83-86), indicating that SEP might have epigenetically induced T cell development (**Fig. 5D**). To confirm this observation, we analyzed the cellular components of the BM of SEP- vs. DMSO-treated groups based on their epigenetic patterns of the genome. We found that the BM of SEP-treated group contained significantly higher total counts of T cells, B cells, and stem cells than DMSO-treated group (**Fig. 5E**). Interestingly, BM-resident T cells are mostly memory T cells returning from distant tissues/organs (87-89), unlike B cells which undergo most of the maturation in the BM (90). Thus, in addition to enhanced lymphogenesis, as indicated by the increases in B cell and stem cell counts, SEP treatment possibly promoted formation of memory T cells to exert long-term protection against tumor formation (91). In fact, certain memory T cell markers, namely, IL6RA, IL7R, IFNGR2, and CD44 (83-87,92), were highly elevated by SEP treatment (**Fig. 5D**), further supporting the possibility of enhanced memory T cell formation by long-term SEP treatment. Our results altogether strongly suggest that SEP might be utilized as a novel immunotherapeutic agent for preventing HER2-positive breast cancer.

**Figure 5.**
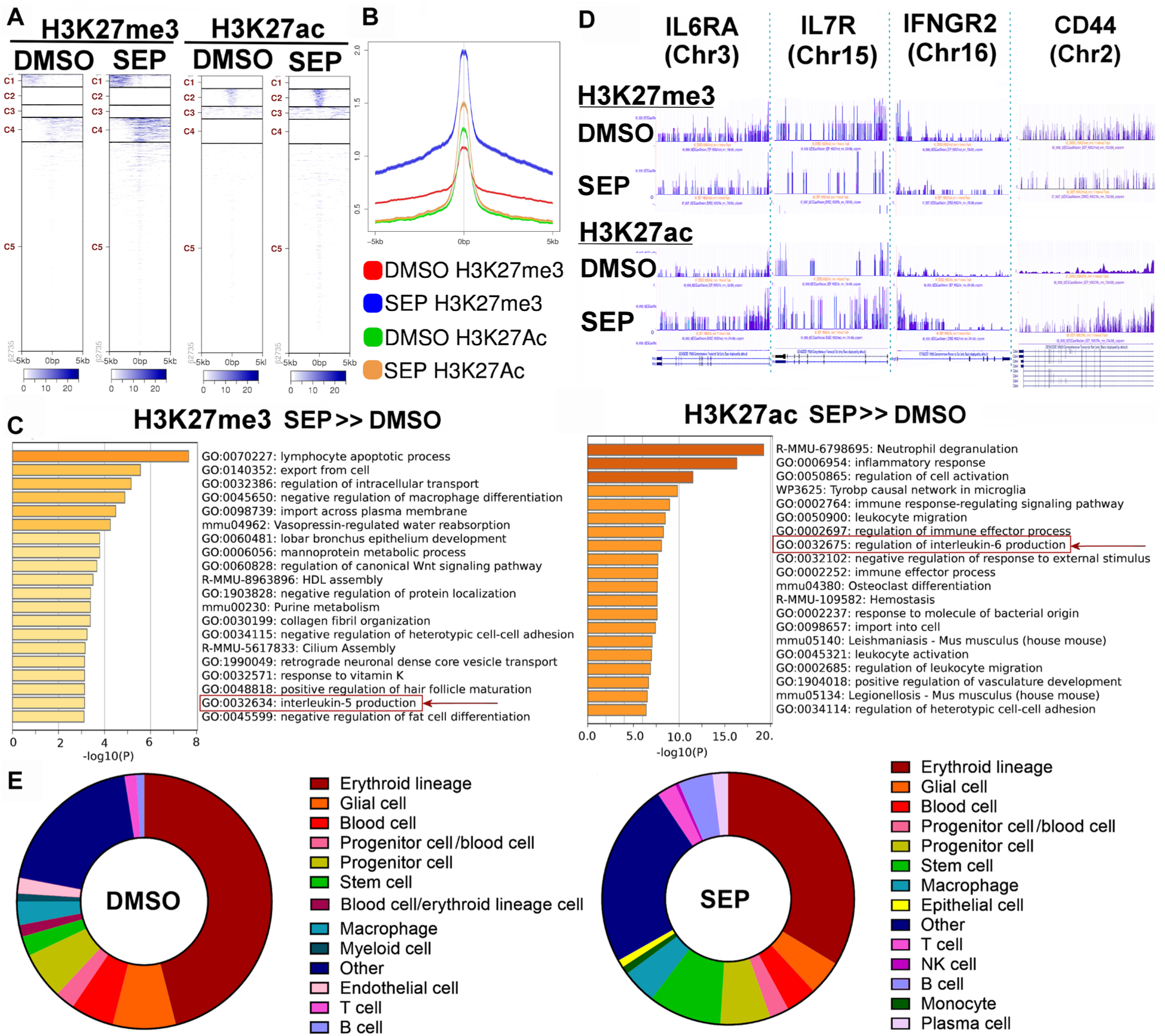
CUT & Tag analysis of bone marrow of SEP-treated MMTV-neu mice reveals increased levels of total T cells. **A-B** The heatmaps (**A**) and peak sizes (**B**) of H3K27me3 (suppression) and H3K27ac (activation) histone marks of the bone marrows of DMSO with tumor vs. SEP without tumor groups of MMTV-neu mice. Note that both histone marks were higher in the latter than the former groups. **C.** Pathways of genes with elevated H3K27me3 mark (more strongly suppressed, left) vs. pathways of genes with elevated H3K27ac mark (more strongly activated, right) in SEP without tumor group compared to DMSO with tumor group. Note that SEP-suppressed pathways belonged to those involved in immune suppression (e.g., IL5 production), whereas SEP-activated pathways belonged to those involved in pro-inflammatory responses (e.g., IL-6 production). **D.** H3K27me3 and H3K27ac marks of select SEP-activated genes involved in T cell activation or memory T cell formation (IL6RA, IL7R, IFNGR2, and CD44 (83-86). **E.** Bone marrow cell compositions determined based on epigenomic profiles (predicted by Cellkb). Note the significant increase in T cell, NK cell and stem cells in the bone marrow of SEP-treated animals.

## Discussion

Recent studies have unveiled dynamic and reciprocal interactions between metabolism and immunity in the body (93). A variety of metabolic signaling pathways not only drive the development of immune cells, but also determine the immunogenicity of tissues/organs and the rest of the body. Conversely, different immunological responses, represented by the expression of unique transcription factors and distinct cytokines, drive cognate metabolic pathways. For example, T helper 1 (Th1) cells (expressing T-bet and IFN-γ) signal to promote glycolysis in macrophages, while downmodulating tricarboxylic acid (TCA) cycle, fatty acid oxidation, and glutaminolysis, inducing their polarization to the pro-inflammatory M1-type. Promotion of glycolysis for energy production is beneficial to M1 TAMs, since they are mostly localized in hypoxic environments that inherently trigger glycolysis (94). In contrast, Th2 cells (expressing GATA3, IL-4, and IL-13) signal to activate peroxisome proliferator-activated receptor gamma (PPARγ) in macrophages to promote lipid metabolism (e.g., fatty acid uptake/oxidation) as well as mitochondrial biogenesis/respiration, inducing their polarization to the anti-inflammatory M2-type (93). Promotion of mitochondrial respiration and fatty acid oxidation for energy production is beneficial to M2-TAMs, since they usually do not have enough glucose supply for glycolysis after its rapid consumption by tumor cells (94). Furthermore, recent studies demonstrated the critical roles of distinct arginine metabolic pathways, namely, NO vs. PAs, in formation and functions of M1- vs. M2-macrophages, respectively (8,19,95,96). Arginine metabolic pathways are also found to play essential roles in regulation of T cell activities. For example, NO promotes T cell proliferation/activation, while elevating the Th1/Th2 ratios (97). Conversely, PAs, elevated in response to T cell activation, help determine T helper cell lineages (Th1, Th2, Th17, and Treg), while inhibiting memory T cell formation (7,98).

Such close linkages between metabolic pathways and immune responses have led to the recent emergence of the concept of Immunometabolism (99,100). A number of metabolic modulators have been developed aiming to target certain metabolic pathways for cancer therapy (101). One of the most explored metabolic pathways is arginine. In cancer, arginine tends to be converted to protumor PAs due to downmodulation of NO synthesis under reduced availability of the NOS cofactor BH_4_ (11,18,19). Different strategies have been explored seeking to normalize arginine metabolism in cancer, where most such efforts have been focused on arginine deprivation therapy and PA synthesis inhibitors (6,13,14,101). Although these approaches have shown some therapeutic benefits, their reported serious adverse side effects (e.g., hearing loss and hematologic disorders) limit their usability (15,16).

In our previous studies, we showed that SEP, the endogenous BH_4_ precursor, could correct arginine metabolism in animal models of HER2-positive mammary tumors. We saw that supplementing SEP induced metabolic and phenotypic reprogramming of tumor cells as well as TAMs and effectively inhibit HER2-positive mammary tumor growth (18,19). Besides, SEP has no dose-limiting toxicity reported during the Phase I trial for phenylketonuria treatment (17). In the present study, we tested the effects of a long-term use of SEP on tumor prevention. We reported that a long-term oral administration of SEP to animals prone to HER2-positive mammary tumors strongly prevented tumor occurrence for 8 months, while most control animals had developed tumors. These SEP-treated animals had undergone the reprogramming of the systemic metabolism and immunity, elevating total T cell counts in the circulation and bone marrow. Given that bone marrow-resident T cells are mostly memory T cells (87-89), it is possible that SEP promoted memory T cell formation, leading to potent tumor prevention. These findings suggest the possible roles of the SEP/BH_4_/NO axis in promoting memory T cell formation, at least in part, by inhibiting PA synthesis known to inhibit CD8+ memory T cell formation (7). Since therapeutic approaches to elevate memory T cells have been yet at the early developmental stages, clinical uses of the SEP/BH_4_ pathway for cancer prevention would warrant further investigation.

## Acknowledgement

We thank Raghvendra Srivastava, Vladimirand Makarov, and Ivan Juric of the Discovery Lab in the Global Center for Immunotherapy and Precision Immuno-Oncology at Cleveland Clinics Foundation for single cell sequencing and analysis. We would also like to thank the research teams at Metabolon, Inc. for metabolomic analyses; Active Motif for CUT&Tag analysis; and Drs. Andrea Kalinoski and David Weaver in the Imaging Core at the University of Toledo for various support in FACS analyses.

## Author Contributions

Conceptualization, V.S, V.F., and S.F; Methodology, V.S, V.F, X.Z., E.C., V.T., O.S., and S.F; Formal Analysis: V.S, V.F, and S.F.; Investigation, V.S., V.F, X.Z., E.C., V.T., and O.S.; Data Curation, V.S, V.F. and S.F; Writing – Original Draft Preparation, S.F.; Visualization, V.S, V.F and S.F.; Supervision, S.F.; Project Administration, S.F.; Funding Acquisition, S.F.

